# Revealing the Paper Mill Iceberg: AI-Based Screening of Cancer Research Publications

**DOI:** 10.1101/2025.08.29.673016

**Authors:** Baptiste Scancar, Jennifer A. Byrne, David Causeur, Adrian G. Barnett

## Abstract

**Objectives:** To train and validate a machine learning model to distinguish paper mill publications from genuine cancer research articles, and to screen the cancer research literature to assess the prevalence of papers that have textual similarities to paper mill papers.

**Design:** Methodological study applying a BERT-based text classification model to article titles and abstracts.

**Setting:** Retracted paper mill publications listed in the Retraction Watch database were used for model training. The cancer research corpus was screened by the model, using the PubMed database restricted to original cancer research articles published between 1999 and 2024.

**Participants:** The model was trained on 2,202 retracted paper mill papers and validated on independent data collected by image integrity experts. A total of 2.6 million cancer research papers were screened.

**Main outcome measures:** Classification performance of the model. Prevalence of papers flagged as similar to retracted paper mill publications with 95% confidence intervals and their distribution over time, by country, publisher, cancer type, research area, and within high-impact journals (Decile 1).

**Results:** The model achieved an accuracy of 0.91. When applied to the cancer research literature, it flagged 9.87% (95% CI 9.83 to 9.90) of papers and revealed a large increase in flagged papers from 1999 to 2024, both across the entire corpus and in the top 10% of journals by impact factor. Over 170,000 papers affiliated with Chinese institutions were flagged, accounting for 35% of Chinese cancer research articles. Most publishers had published substantial numbers of flagged papers. Flagged papers were overrepresented in fundamental research and in gastric, bone, and liver cancer.

**Conclusions:** Paper mills are a large and growing problem in the cancer literature and are not restricted to low impact journals. Collective awareness and action will be crucial to address the problem of paper mill publications.

## Introduction

Research paper mills are ‘*contract-cheating organisations which provide undeclared services to support research manuscripts and publications*’^1^. Research paper mills fabricate and submit manuscripts for their customers. The first research paper mill activity was reported in the 2010s^2^, and has since increased in volume and sophistication. More than 400,000 papers suspected to have originated from paper mills have been published in the last 20 years^3^, with the paper mills earning tens of millions of dollars annually^4^. This issue gained visibility when Wiley – after acquiring Hindawi – retracted nearly 11,000 suspected paper mill papers and shut down 19 journals over two years^5^.

Research paper mills maximise their earnings by quickly producing industrial quantities of low quality research papers^6,7^. To produce manuscripts at scale, fabrication has likely relied on templates with pre-made sentences where domain-specific terms vary^8^. Suspect papers can include incorrect reagents^9^, fabricated data and experiments^2,10^, and photoshopped or re-used figures^11^. Paper mill papers are often generic, poorly written, lack coherence between sections^8,12^, and may offer superficial research justifications^7^. Paper mills sell manuscripts to researchers eager to increase their number of publications^10^, creating author groups who have never worked together or made any intellectual input^13^. Paper mills often cite their own productions^14^, possibly as part of their service to clients who pay for citations^15^. Paper mills may even bribe editors and manipulate peer-review to facilitate publication, as shown by online discussions between researchers and likely paper mill contacts^4^.

Paper mill manuscripts can be simultaneously submitted to multiple journals until acceptance, wasting the time of editors and reviewers^2,11,16^. The reported percentage of paper mill submissions to journals ranges from 2 to 46%^2^. Paper mills likely target journals where fabricated manuscripts have already been accepted^2^, increasing their chances of success. They may also focus on high-impact journals, as the prices they charge – according to A. Abalkina’s investigations of a paper mill^16^ – can be directly linked to the journal’s impact factor^17^.

The overall prevalence of paper mill papers in biology and medicine is estimated as 3%^3^ but cancer research and particularly molecular oncology could be more affected^1^. This can be explained by high publication pressure^7^, a specialised field with simple-to-fake data and techniques^1^, and limited peer-review capacity^6^ – making fake papers easier to produce and harder to detect.

The rise of AI could exacerbate the paper mill problem, via automated image and text generation, making detection more challenging^18^. Some publishers are using screening tools to detect manuscripts from paper mills^19,20^. Integrity sleuths have also developed detection methods, such as identifying awkward rewording of scientific terms, known as ‘*tortured phrases*’^21^, or nucleotide sequence reagent verification^9^. Paper mill manuscripts may have missing or unusual acknowledgements, funding or ethics statements^11^.

Previous publications have documented recurrent formatting templates among suspected paper mill papers, including identical layouts and nearly identical textual formulations in figure legends^8,9,13,22,23^. Bless *et al*.^24^ have reported ‘*distinct phrase patterns and higher word repetition*’ and ‘*lower lexical diversity*’ in full texts and abstracts of retracted papers. Also, Bless *et al.*^24^ and other studies^25,26^ have demonstrated that machine learning methods can predict retractions and paper mill products from text using data from Retraction Watch – a non-profit group that records paper retractions^27^ – in a cross-disciplinary context rather than within a specific scientific domain.

The performance of such an approach has never been tested in cancer research, and we believe it has the potential to systematically screen papers to assess the prevalence of papers sharing textual similarities with paper mill publications. We hypothesise that paper mills use text templates, which likely extend to titles and abstracts. We also believe that these templates are domain- and publication type-specific and could constitute strong signals to machine learning models. Thus, we aim to use cancer research titles and abstracts from retracted paper mill papers as input to a BERT-based (Bidirectional Encoder Representations from Transformers)^28^ machine learning pipeline for a text classification task. BERT learns from examples to recognise patterns in text, enabling it to identify when new papers share similarities with retracted paper mill papers. Using only titles and abstracts enables easy access to training data and offers scalability for producing broad estimates.

The first objective was to train and evaluate our model’s ability to reliably classify titles and abstracts from retracted papers attributed to suspected paper mill activity and genuine cancer research papers. The second objective was to use our model to screen millions of cancer research papers to assess the prevalence of flagged papers over time, across countries, publishers, and cancer research subdomains, and to examine how they have evolved in high– impact factor journals (defined as Decile 1, or D1 – top 10% journals in this study). Throughout, we use flagged papers to refer to articles whose titles and abstracts are textually similar to retracted papers tagged as ‘*paper mill*’ in Retraction Watch. Flagging is a statistical screen, not an attribution of misconduct.

This research aims not only to assess the potential of machine learning for detecting cancer research papers that share textual features with paper mill manuscripts, but also to raise awareness of potentially fabricated papers among stakeholders in cancer research. We believe that such research could support editorial triage, help protect genuine researchers from citing or using fake research, inform funding and institutional policies on research integrity, and provide insights into the scale and dynamics of paper mill activity within cancer research.

## Material and methods

### Cancer research corpus

#### PubMed baseline pre-processing

To create a comprehensive cancer research dataset, the entire biomedical research corpus from the 2025 PubMed (https://pubmed.ncbi.nlm.nih.gov/) database was downloaded in March 2025. The following data were extracted from each of the over 38 million papers: PubMed Identifier (PMID), title, abstract, original language, journal name, journal ISSN (International Standard Serial Number), publication date, first author’s affiliation, publication type and MeSH terms (Medical Subject Headings). The data were pre-processed following the method used by González-Márquez *et al.*^29^ as follows: we excluded abstracts that were non-English, empty, truncated and unpunctuated to avoid these features influencing the language model. The text was transformed into standardised tokens (units of text such as words or punctuation marks) for analysis. Abstracts of less than 250 tokens and more than 4,000 tokens were removed as these were generally non-standard abstracts and were also rare (<1%). After this initial filtering, the research dataset contained 24.8 million papers published between 1975 and 2025.

Next, the dataset was filtered to retain only papers published within the target time frame (1999 to 2024): papers prior to 1999 or after 2024 were excluded, and duplicates were removed, reducing the dataset to 20.2 million papers. We only included papers classified as ‘*Journal Article*’ and excluded other publication types, such as literature reviews and clinical trials. These types may also be targeted by paper mills, but would require separate models, as we expect paper mills to use manuscript type-specific templates. All retraction notices, corrections and expressions of concern were also removed. After this second filtering step, 17.4 million papers remained.

#### Cancer research filtering

The cancer research corpus was derived from the remaining papers using a two-level keyword filtering strategy. *Box 1* shows the keywords searched for in titles and abstracts of the 17.4 million papers. These keywords were adapted from MeSH terms and National Cancer Institute^30^ terminology. The keyword matching was designed to be specific to cancer whilst also retaining the broadest coverage of cancer research papers. We acknowledge that some non-cancer-related papers may be included and that not all cancer-specific terms have been used.

#### Box 1: Cancer-related keywords

astrocytoma, carcinoembryonic antigen, carcinoid, carcinogen, carcinogenesis, carcinoma, cancer, checkpoint inhibitor, chemotherapy, chordoma, ependymoma, glioblastoma, glioma, leukaemia, leukemia, lymphoma, macroglobulinaemia, macroglobulinemia, medulloblastoma, melanoma, mesothelioma, metastasis, metastatic, myelodysplastic syndrome, myeloma, myeloproliferative neoplasm, neuroblastoma, nsclc, oncogene, oncogenesis, oncology, pheochromocytoma, radiation therapy, radiotherapy, retinoblastoma, sarcoma, seminoma, tumor and tumour.

*Substring matching* ensured that terms such as *osteosarcoma* were captured under broader categories like *sarcoma*. Papers matching multiple keywords were included only once to avoid duplicates. Both UK and US spellings of each cancer-related term were used. All levels of the search were combined to produce a final cancer research dataset of 2,647,471 papers, published across 11,632 journals. A flowchart detailing the filtering strategy from the initial PubMed database to the final cancer research corpus is provided in *Figure 1*.

**Figure 1:**
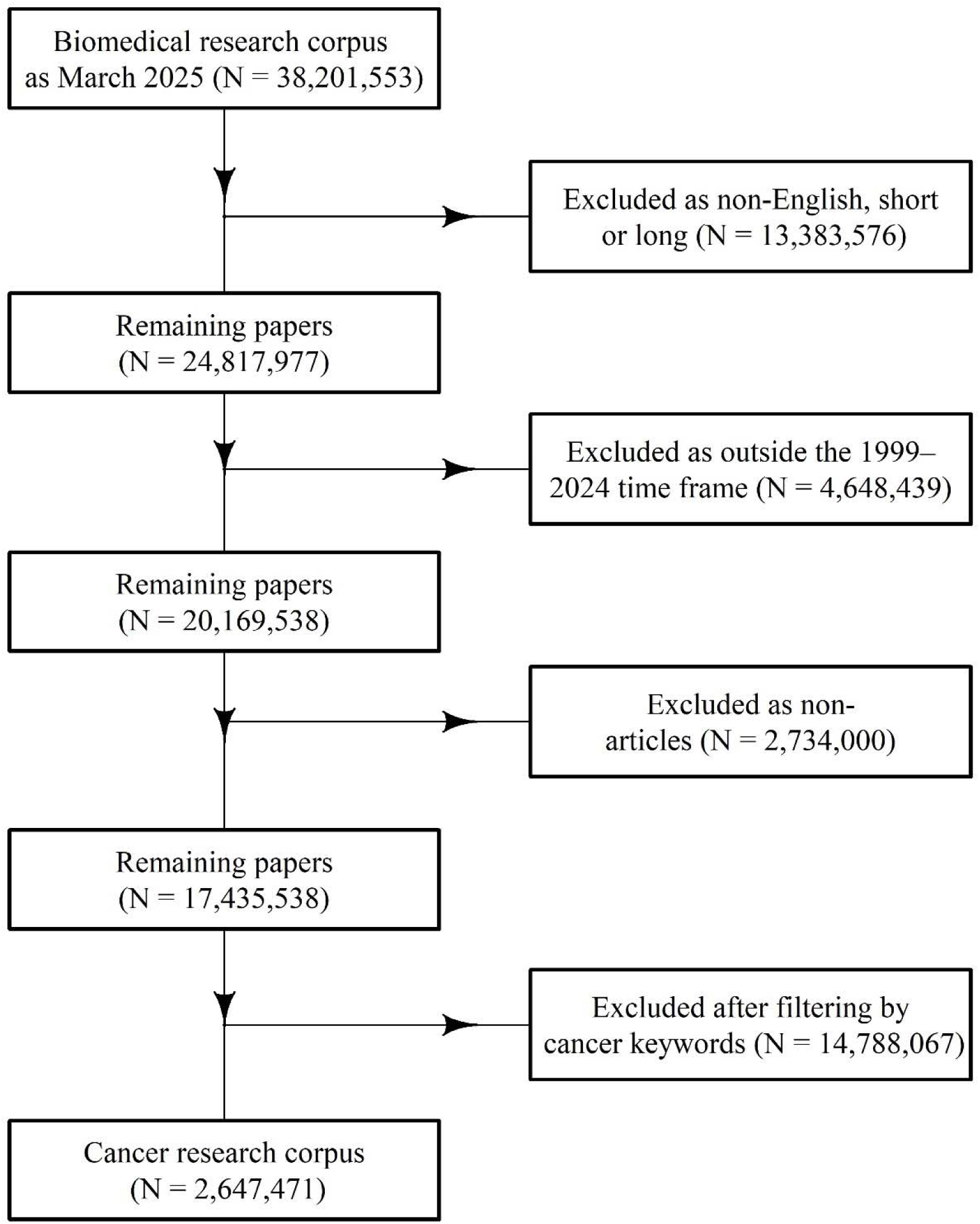
Flow diagram showing all preprocessing steps and exclusions applied to derive the final cancer research corpus. Cancer keywords are provided in Box 1.

Data extracted for visualisation purposes are the first author’s country of affiliation, the publisher, the type of cancer investigated, the main cancer research areas, and the SCImago Journal Impact Factor^31^. The type of cancer investigated and the main cancer research areas were inferred from the titles, abstracts, and MeSH terms of the papers using AI-based labelling. All extraction methods are detailed in *Supplementary File 1.* A detailed breakdown of these variables describing the cancer research corpus is provided in *Supplementary File 2*.

### Paper mill datasets

We developed our model using two sources of paper mill papers: 1) Papers tagged as originating from paper mills in the Retraction Watch Database^27^; 2) Online lists compiled by research image integrity experts – also called integrity sleuths – where evidence of image manipulation was found. A compilation of paper mill papers is available online in the ‘*Spreadsheet of spreadsheet’* thanks to anonymous PubPeer contributors^32^. PubPeer^33^ is a web site (https://pubpeer.com/) that allows users to leave post-publication comments concerning potential research integrity issues or other concerns. It has been used in research on integrity^34^ and has played an important role in high-profile retractions^35^.

The Retraction Watch dataset was used for model development (training, optimisation and internal validation), whereas the experts’ dataset was reserved for external validation of model performance. From the 64,457 total retractions recorded in the Retraction Watch Database as of June 2025, we removed papers that were not in the PubMed database, and those that were not retractions (e.g., Expressions of Concern, Corrections, or Reinstatements). The ‘*Paper Mill*’ tag in the retraction reason field was used to identify 5,657 retracted publications. Only those whose PMIDs matched entries in our cancer research corpus were retained, reducing the number to 2,270 retracted papers. We excluded papers for which the original text had been replaced by the retraction notice, reducing the number of retracted paper mill papers to 2,202.

To validate the model’s ability in new data, we used 3,094 suspected paper mill papers from the integrity experts’ dataset for external model validation, excluding those that overlapped with the Retraction Watch set. These papers were also extracted from the cancer research corpus.

Visualisations of data used at the training stage from the Retraction Watch and the integrity experts’ sets are presented in *Supplementary File 3*. These show the distribution of publishers, countries of the first authors’ institutions, cancer types, and research areas among paper mill papers, as well as the single word, two-word and three-word combinations most prevalent in the titles of paper mill papers.

### Model selection and training

#### Controls selection

Our training dataset was chosen to be balanced, with 50% paper mill papers and 50% presumed genuine papers. Controls were selected from the cancer research corpus with the aim of including as few paper mill papers as possible to minimise bias and enhance training performance. Given the difficulty of assessing the genuine status of cancer research papers in large samples, control papers were selected from high impact factor journals (Decile 1) and countries underrepresented in the Retraction Watch database (using the country of the first author’s institution). Throughout this paper, controls should be regarded as proxies for quality research, not verified genuine papers.

To avoid a potential bias in the English diction used by Chinese authors, we included 101 papers (5%) in the control set authored by researchers from Chinese institutions that were published in four high-impact journals: *Cell*, *Cancer Cell*, *Molecular Cell*, and *The EMBO Journal*. We also included 600 of the most cited Taiwanese cancer research papers (28%) listed in OpenAlex^36^, as Taiwan is a predominantly Mandarin-speaking country, but is less represented in the Retraction Watch database (with only one recorded retraction due to paper mill involvement in the cancer research corpus). Another 33% of the control papers were randomly selected from Swedish, Finnish and Norwegian institutions in cancer research, as these countries have no recorded instances of paper mill retractions in the Retraction Watch database. The remaining 33% consisted of a random selection of papers published in *Cell*, *Cancer Cell*, *Molecular Cell*, and *The EMBO Journal*, from countries other than China, Taiwan, Norway, Sweden and Finland.

For the external validation dataset, papers from the integrity experts’ dataset were combined with a similar number of control papers, randomly sampled from authors from Swedish, Finnish and Norwegian institutions, and other papers published in the four high-impact factor journals mentioned above. None of the controls or paper mill papers used in the training set overlap with those in the external validation set.

All papers in the control sets were manually verified as free of research integrity concerns on PubPeer (June 2025). A paper was removed if any problem was reported. We assumed that all papers in the paper mill datasets are indeed paper mill products, and that the controls are legitimate scientific papers. However, we acknowledge that the ground truth is unknown and that, despite our efforts, some papers may have been mislabelled.

#### Model selection and training

We chose to use only titles and abstracts to train the model, as these data were often available (full texts are frequently behind paywalls). Each paper’s title and abstract were combined. We framed the detection of paper mill papers as a binary text classification problem – either *Authentic* or *Fraudulent* – to provide supervision signals to the model.

The 4,404 labelled papers from the Retraction Watch dataset and its controls (2,202 paper mill papers and 2,202 controls) were split into 70% for training (n = 3,082), 17.5% for optimisation (n = 771), and 12.5% for internal validation (n = 551). The model was trained and tuned using the training and optimisation sets. Splitting was performed at the paper level and stratified by class label to preserve the 1:1 ratio of paper mill and control papers across all subsets.

Model performance (accuracy, sensitivity and specificity) was first evaluated on the internal validation set – which was never seen during training or optimisation – and then on an external validation set composed of papers from research integrity experts (3,094 paper mill papers and 3,100 controls). The latter dataset was entirely independent from Retraction Watch and used to assess the model’s generalisation to new data. A summary of all datasets and their respective roles is provided in *Table 1*. Each metric was derived from the confusion matrix using a probability threshold to prioritise specificity and minimise false positives. Reported values correspond to a single final model obtained after hyperparameter optimisation over 150 trials. Confidence intervals were not computed, as performance was estimated on fixed validation sets. This is further detailed in *Supplementary File 1*.

**Table 1:**
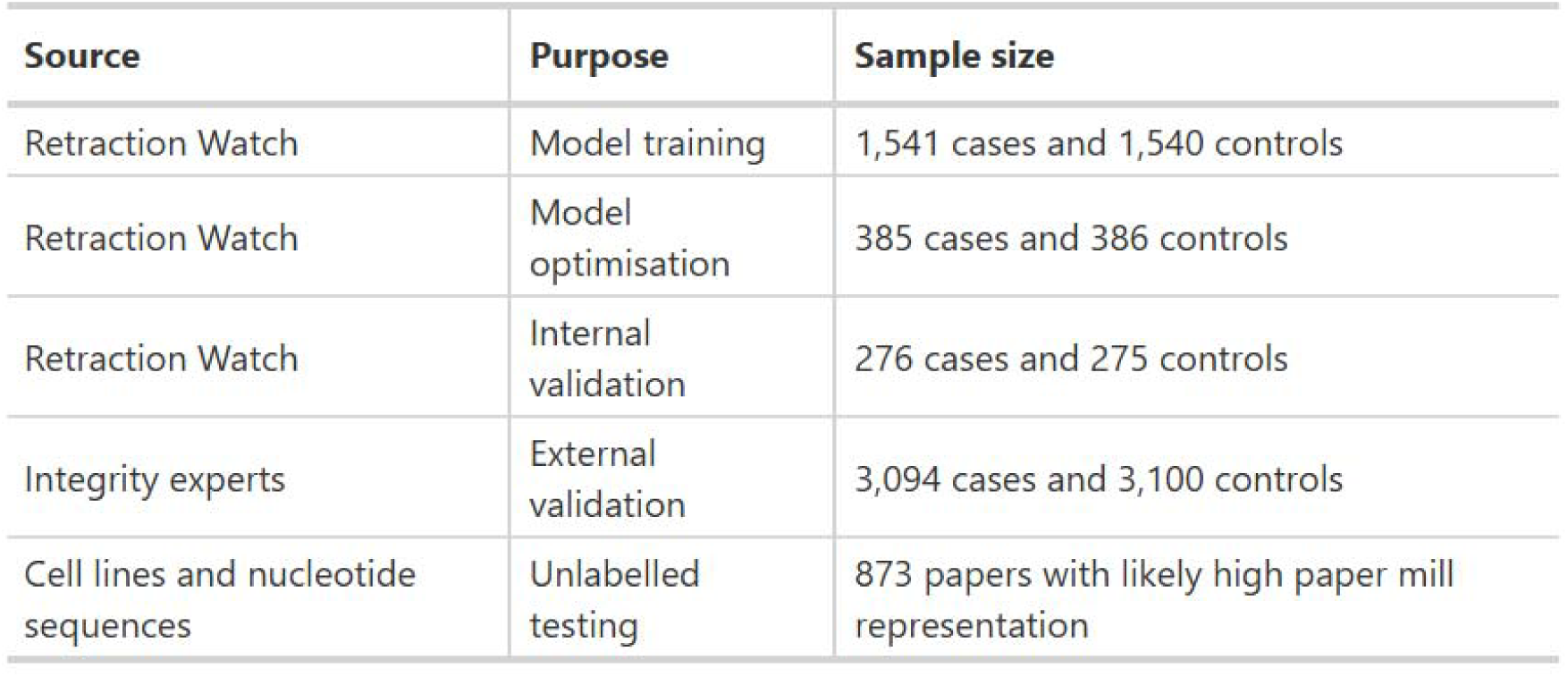
summary of the five datasets and their use in the experiment design.

We selected BERT to analyse the text for its strong performance and relatively low computational cost, due to its moderate number of parameters^28^. The model was initialised from the public BERT-base-uncased checkpoint and fine-tuned on our dataset with a new classification head. The model was initially compared with RoBERTa, BioBERT, PubMedBERT, Longformer, and Clinical Longformer^37–41^ to assess the potential of biomedical-specific pretraining and extended input capacity. Details of comparative model testing are provided in *Supplementary File 4*.

All journal-specific formatting was removed from the abstracts. Because BERT truncates inputs beyond its 512-token limit, each title and abstract was split into individual sentences to prevent information loss. Each sentence was labelled (paper mill or not) and fed into BERT during the training phase. During inference, predictions were made at the sentence level and final classification probabilities were obtained by averaging the positive class probabilities across the title and abstract. The optimisation method is described in *Supplementary File 1*. A logistic regression model was also tested on BERT’s output probabilities to incorporate sentence ordering; however, it did not outperform the simple averaging approach and so was not used (*Supplementary File 1*).

#### Additional validation

To further evaluate the model, we checked whether 873 problematic cancer research papers reported in three prior studies where paper mill involvement was suspected ^42–44^, were flagged by the model (*Table 1*). These papers were not necessarily from paper mills, as similar errors can sometimes occur in genuine research. This analysis was not intended to measure model performance directly but rather to assess the model’s ability to flag papers that had already been reported as suspicious. While we did not expect all these papers to be flagged, we did anticipate that most would be.

None of these papers were included in the training or validation sets. All papers overlapping with paper mill papers or controls from the training, optimisation or validation sets were removed during data extraction (n = 29). No content re-evaluation was performed, and integrity-related information was not disclosed to the model. These papers, involving misidentified and/or non-verifiable nucleotide sequences (primers and other oligonucleotides used for amplification, detection, or gene knockdown) or cell lines, were retrieved from three sources: 193 cancer-related papers identified by Oste *et al*.^42^, 113 cancer-related papers in high impact factor journals – *Molecular Cancer* and *Oncogene* – reported by Pathmendra *et al*.^43^, and 567 cancer-related papers, as listed by Park *et al*.^44^.

#### Patient and Public Involvement

There was no involvement in this research of patients or the public.

### Screening the cancer literature

Each of the 2.6 million papers of the cancer research corpus published between 1999 and 2024 was screened with our fine-tuned version of BERT trained to identify textual features associated with retracted paper mill papers. We refer to papers that were classified as resembling retracted paper mill papers as ‘*flagged papers*’. None of the papers seen by the model during training were excluded from the cancer research corpus (n = 3,853). These papers were considered part of the corpus and represented a small percentage of flagged papers and so not large enough to influence the overall conclusions.

The 95% confidence interval for the proportion of flagged papers was estimated via bootstrapping (1,000 resamples with replacement). All figures are presented as percentages of the total number of papers, to illustrate the proportion of non-flagged papers compared to flagged papers. Data were visualised using the ggplot2 R library^45^. A diagram of the study design is in *Figure 2*.

**Figure 2:**
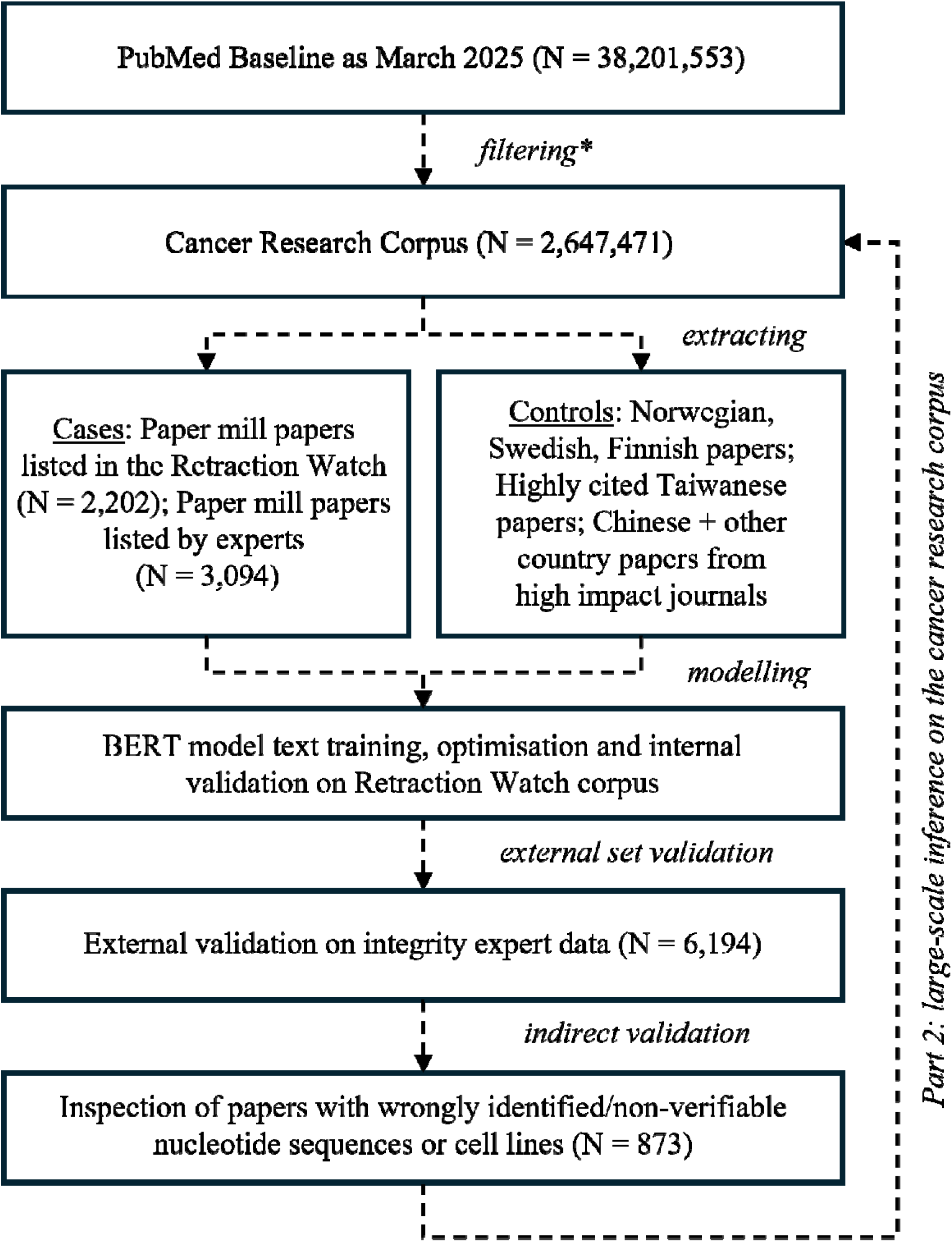
Study design and workflow for large-scale detection of papers textually similar to paper mill papers in the cancer research corpus. Filtering method (*) is shown in Figure 1.

## Results

### Model performance and errors

#### Training data

Retraction Watch covered papers from 2007 to 2024, with a peak in 2022, while the integrity experts’ data included papers from 2010 to 2024, with a peak between 2019 and 2020 (*Figure S3.1*). In the Retraction Watch dataset, the most frequent publisher is John Wiley & Sons (via Hindawi), followed by Spandidos Publications and Informa (*Table S3.2*). In the experts’ dataset, Springer Nature has the most suspected paper mill papers, followed by John Wiley & Sons and Spandidos Publications (*Table S3.4*). Papers from Chinese institutions are highly represented in both datasets (*Tables S3.2 and S3.4*). Common topics in paper titles for both datasets include microRNAs, long non-coding RNAs, and lung cancer-related terms (*Tables S3.1 and S3.3*). Many papers cover topics related to gene and preclinical research. The most frequent cancer types are lung, liver, colorectal, gastric, and brain cancer (*Tables S3.2 and S3.4*). The most prevalent research fields are *cancer biology* and *cancer therapy*, followed by *cancer diagnosis* (*Tables S3.2 and S3.4*).

#### Performance

Model performance metrics for both datasets are summarised in *Table 2*. The classification model achieved an accuracy of 0.91 in detecting paper mill papers in the Retraction Watch dataset (502 out of 551 papers were correctly classified as either paper mill-associated or genuine), with a sensitivity of 0.87 (239 out of 276 paper mill papers correctly identified) and a specificity of 0.96 (263 out of 275 genuine papers correctly identified). The titles and abstracts of three retracted paper mill papers, all correctly predicted with high probability, are presented as examples in *Supplementary File 5*.

**Table 2:** Model performance over internal and external validation sets.

| Set | Accuracy | Sensitivity | Specificity |
| --- | --- | --- | --- |
| Internal validation | 91% (502/551) | 87% (239/276) | 96% (263/275) |
| External validation | 93% (5,771/6,194) | 87% (2,698/3,094) | 99% (3,073/3,100) |

In validating on unseen data for the integrity experts’ dataset, the model had a similar classification accuracy of 0.93 (5,771 out of 6,194) with a sensitivity of 0.87 (2,698 out of 3,094) and a specificity of 0.99 (3,073 out of 3,100).

In further validation sets, the model flagged 67% (130 out of 193) of problematic papers found by Oste *et al*.^42^, 75% (425 out of 567) of problematic papers found by Park *et al*.^44^, and 66% (75 out of 113) of problematic papers found by Pathmendra *et al*.^43^. Approximately 72% of the combined problematic papers were flagged by the model.

#### Misclassifications

We combined the misclassifications from both the internal and external validation sets, as their characteristics were similar (*Supplementary File 3*). Pooling these data increased the sample size and therefore improved the robustness of analyses. False positives – control papers incorrectly predicted to resemble paper mill papers – were rare, with only 39 out of 3,375 across both the Retraction Watch and integrity experts’ datasets. This small number of false positives did not allow for the identification of generalisable patterns. In contrast, 433 out of 3,370 paper mill papers were false negatives – paper mill papers incorrectly predicted as authentic.

The main characteristics of false negative predictions by the model are summarised in *Table 3* and an extended analysis is *in Supplementary File 6*. False negatives are disproportionately associated with first authors affiliated in China as they account for 90% of false negatives (compared with 47% in the overall validation dataset). Papers published in 2014 (+ 9%) and 2015 (+8%) were more often misclassified as false negatives by the model. Gastric cancer (+4%), liver (+4%), colorectal (+4%), and lung cancer (+4%), as well as publishers such as Rapamycin Press LLC (+4%) and John Wiley & Sons (+2%) are slightly overrepresented among the false negatives. Epidemiology papers are slightly overrepresented among false negatives, while text analyses do not show overrepresented topics other than general cancer words in the false negatives (*Supplementary File 6*).

**Table 3.** Characteristics of false negatives (n = 433) by publication year, publisher, first author’s country and cancer type. For each variable, the 10 most frequent categories were retained. A single Pearson’s chi-squared test of independence was carried out for each variable. The table lists categories that were significantly overrepresented in each variable, defined by standardised residuals greater than 2 (this is not tested and is only descriptive). The percentages are shown both within the false negatives and in the overall pooled validation dataset (n = 6,745). The difference represents false negatives minus overall.

| Variables | Most Overrepresented Categories (residuals >2) | Percent in False Negatives (%) | Percent in Overall (%) | Diff (%) | $\chi^2$ p-value |
| --- | --- | --- | --- | --- | --- |
| Year | 2014 | 12% | 3% | 9% | < 0.001*** |
|  | 2015 | 13% | 5% | 8% |  |
| Publisher | Rapamycin Press LLC | 7% | 3% | 4% | < 0.001*** |
|  | John Wiley & Sons | 14% | 12% | 2% |  |
| Country | China | 90% | 47% | 43% | < 0.001*** |
| Cancer type | gastric cancer | 8% | 4% | 4% | < 0.001*** |
|  | liver cancer | 10% | 6% | 4% |  |
|  | colorectal cancer | 12% | 8% | 4% |  |
|  | lung cancer | 12% | 8% | 4% |  |

### Flagged papers in the cancer literature

After applying our model to the cancer research corpus from 1999 to 2024, there were 261,245 papers flagged as including textual characteristics of retracted paper mill papers – representing 9.87% (95% CI 9.83 to 9.90) of all original cancer research papers.

#### Trends in flagged papers

The number of flagged papers had a clear and rapid increase between 1999 and 2022 (*Figure 3*), peaking in 2022 before showing a slight decline in 2023 and 2024. The number of flagged papers per year followed an exponential trend from 1999 to 2022 (R² for exponential fit = 0.92). While the percentages of flagged papers remained around 1% in the early 2000s, they progressively rose to exceed 15% of the annual cancer research output by the early 2020s.

**Figure 3:**
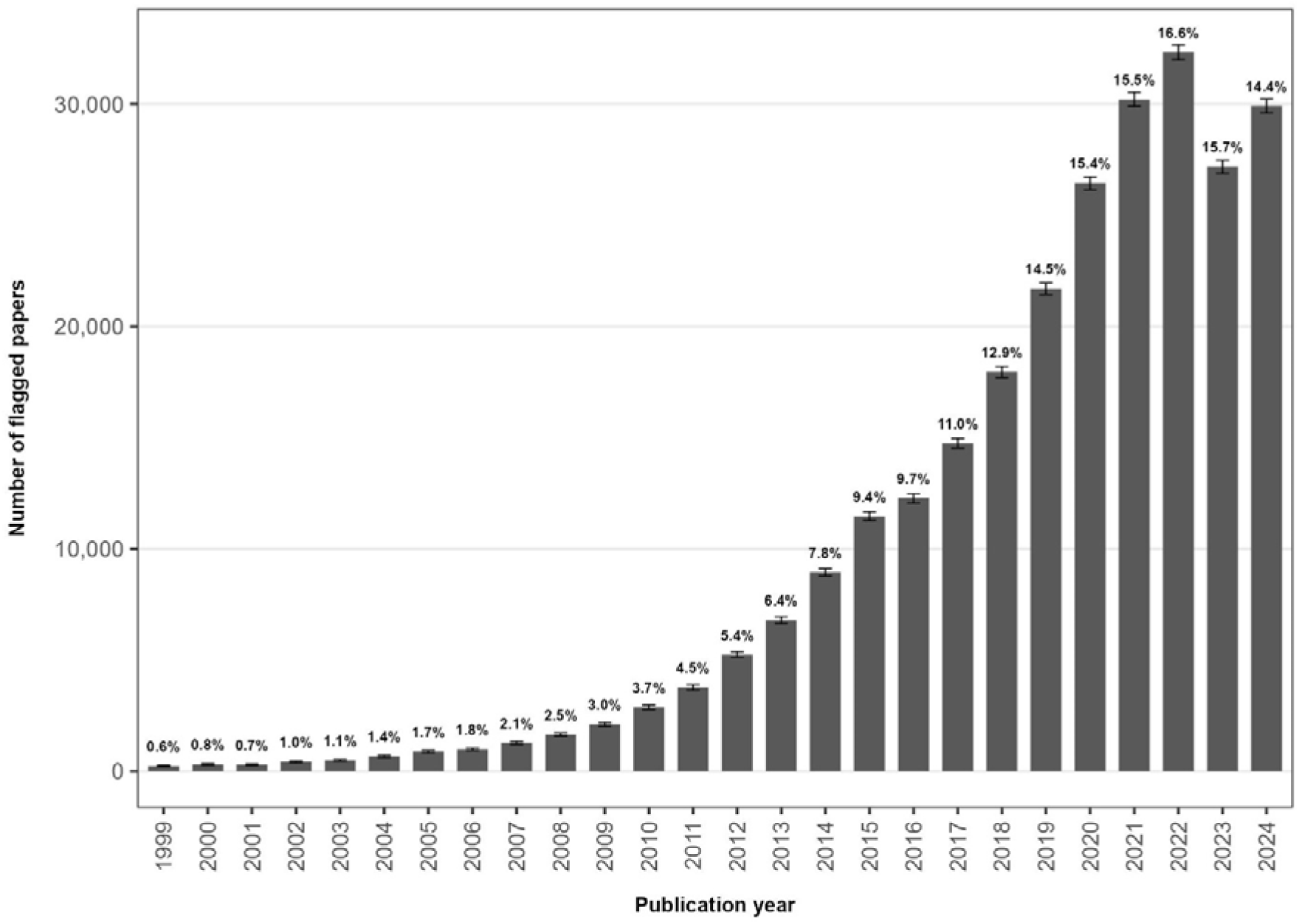
Number of papers per year in the cancer research corpus, flagged because their titles and abstracts were similar to those of retracted paper mill papers. The percentage of flagged papers among all cancer research papers published each year is shown above the bars. Error bars are 95% confidence intervals estimated using bootstrap resampling.

#### Countries of flagged papers

The percentages of flagged papers per country show that papers from China were most frequently flagged, representing 35% of cancer papers with 177,907 flagged papers (*Figure 4*), followed by Iran with 20% of flagged papers, accounting for 6,801 papers. Papers from four other countries were also frequently flagged: Saudi Arabia (16%), Egypt (15%), Pakistan (14%) and Malaysia (13%). The United States was the second country in terms of numbers of flagged papers, with 10,511 flagged papers, representing 2% of US cancer research papers.

**Figure 4:**
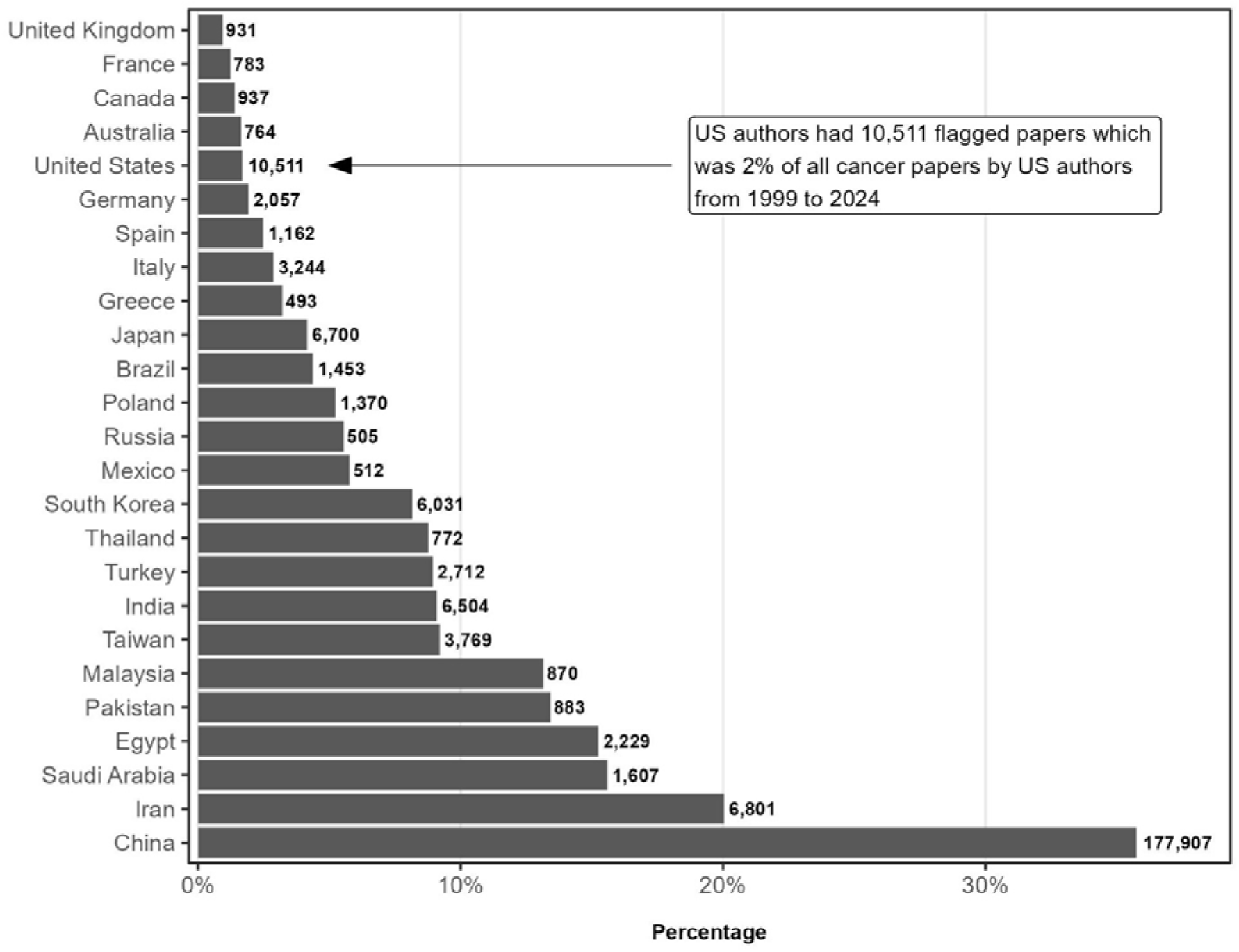
Percentage of papers in the cancer research corpus, flagged by our model because their titles and abstracts were similar to those of retracted paper mill papers, across the 25 countries with the highest numbers of flagged papers based on the first author’s affiliation country. The numbers of flagged cancer research papers per country are shown next to each bar.

#### Publishers and journals of flagged papers

The publisher Verduci Editore had the highest percentage of flagged papers with approximately 65% in its cancer research journal – *The European Review for Medical and Pharmacological Sciences* (*Figure 5*). The second publisher in terms of percentage was International Scientific Literature, with approximately 45% of papers flagged in one journal, *Medical Science Monitor (MSM)*. The next five publishers were E-Century Publishing Corporation (42%), Spandidos Publications (37%), Ivyspring International Publisher (31%), and IOS Press (30%).

**Figure 5:**
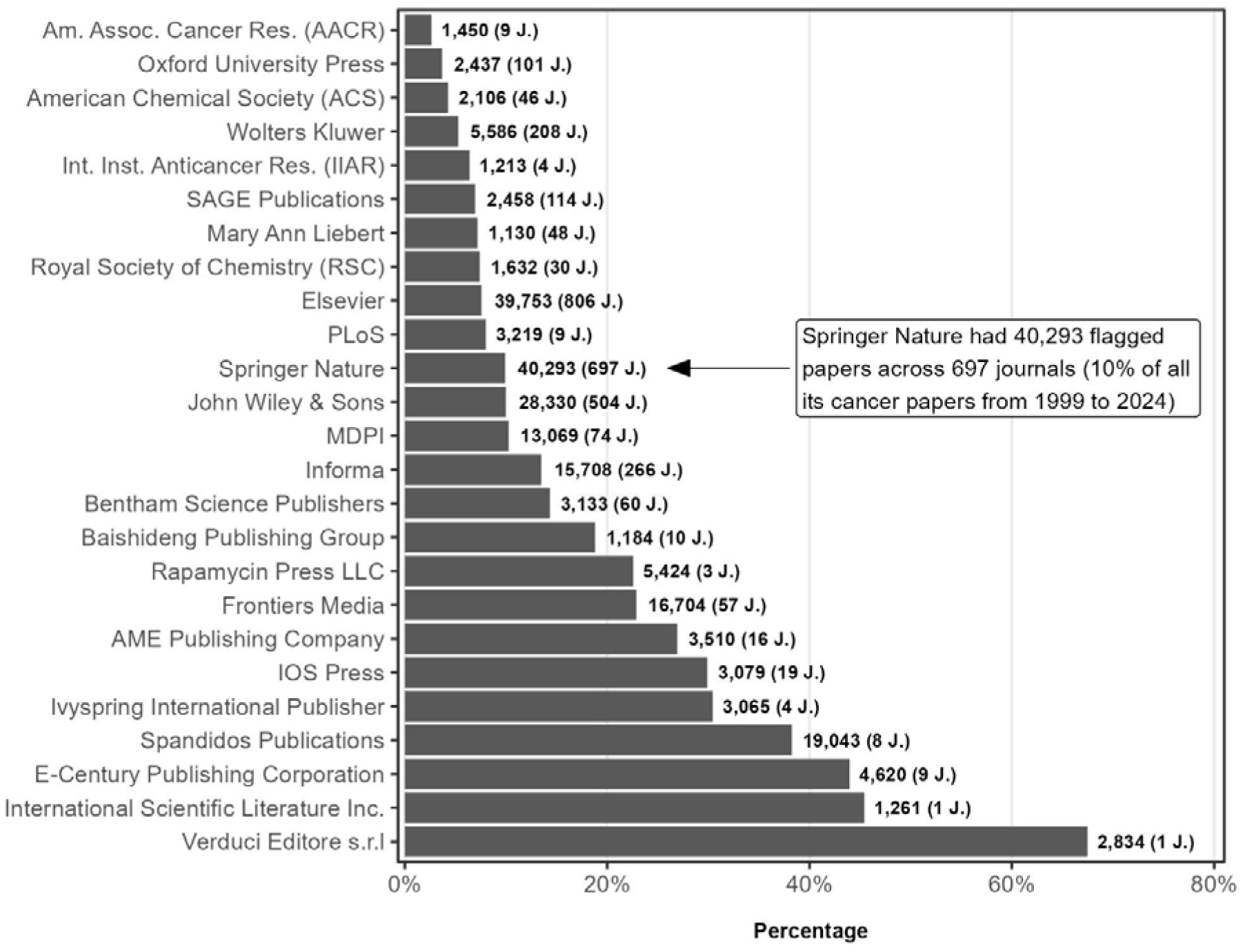
Percentage of papers in the cancer research corpus, flagged because their titles and abstracts were similar to those of retracted paper mill papers, for the 25 publishers with the highest numbers of flagged papers. The number of flagged papers per publisher is given next to each bar and the corresponding number of journals is shown in parentheses.

The largest publishers – such as Elsevier, Springer Nature, and John Wiley & Sons – have a relatively low percentage of flagged papers (around 10%) but account for the highest absolute numbers of flagged papers across more than 500 journals, with flagged paper numbers of 39,753 (Elsevier), 40,293 (Springer Nature), and 28,330 (John Wiley & Sons).

#### Cancer types of flagged papers

Among all cancer types, gastric cancer papers show the highest percentage of flagged papers, with 22% of these papers flagged (*Figure 6*). Bone cancers, such as osteosarcoma, follow with 21% of papers flagged, followed by liver cancer at 19%. Most cancer types fall within a range of 10 to 15% flagged papers. Breast, skin, prostate, and blood cancers show the lowest percentages of flagged papers. In terms of absolute numbers, lung (28,435) and liver (26,730) cancer account for the highest numbers of flagged papers.

**Figure 6:**
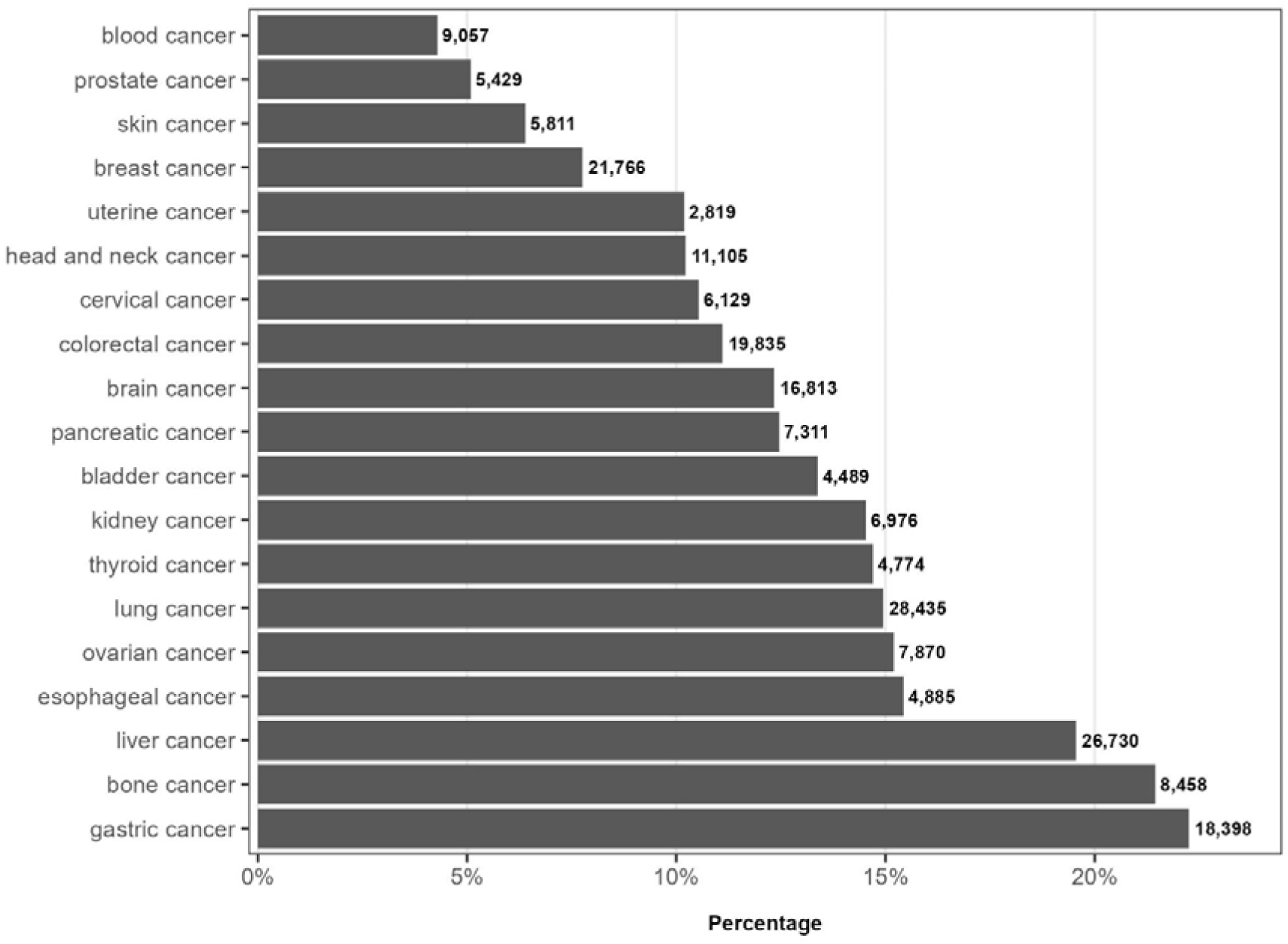
Percentage of cancer research papers according to cancer type, flagged because their titles and abstracts were similar to those of retracted paper mill papers. The number of flagged papers per cancer type is shown next to the bars.

#### Cancer research area of flagged papers

Flagged papers are largely concentrated in *Cancer Biology and Fundamental Research*, as well as in *Treatment Development or Evaluation* and *Diagnosis and Prognosis*, where percentages exceed 10% (*Figure 7*). In contrast, areas such as *Survivorship, Supportive Care and end-of-life*, *Epidemiology and population studies*, and *Health Systems, Policy and Implementation* had lower percentages of flagged papers (under 2%).

**Figure 7:**
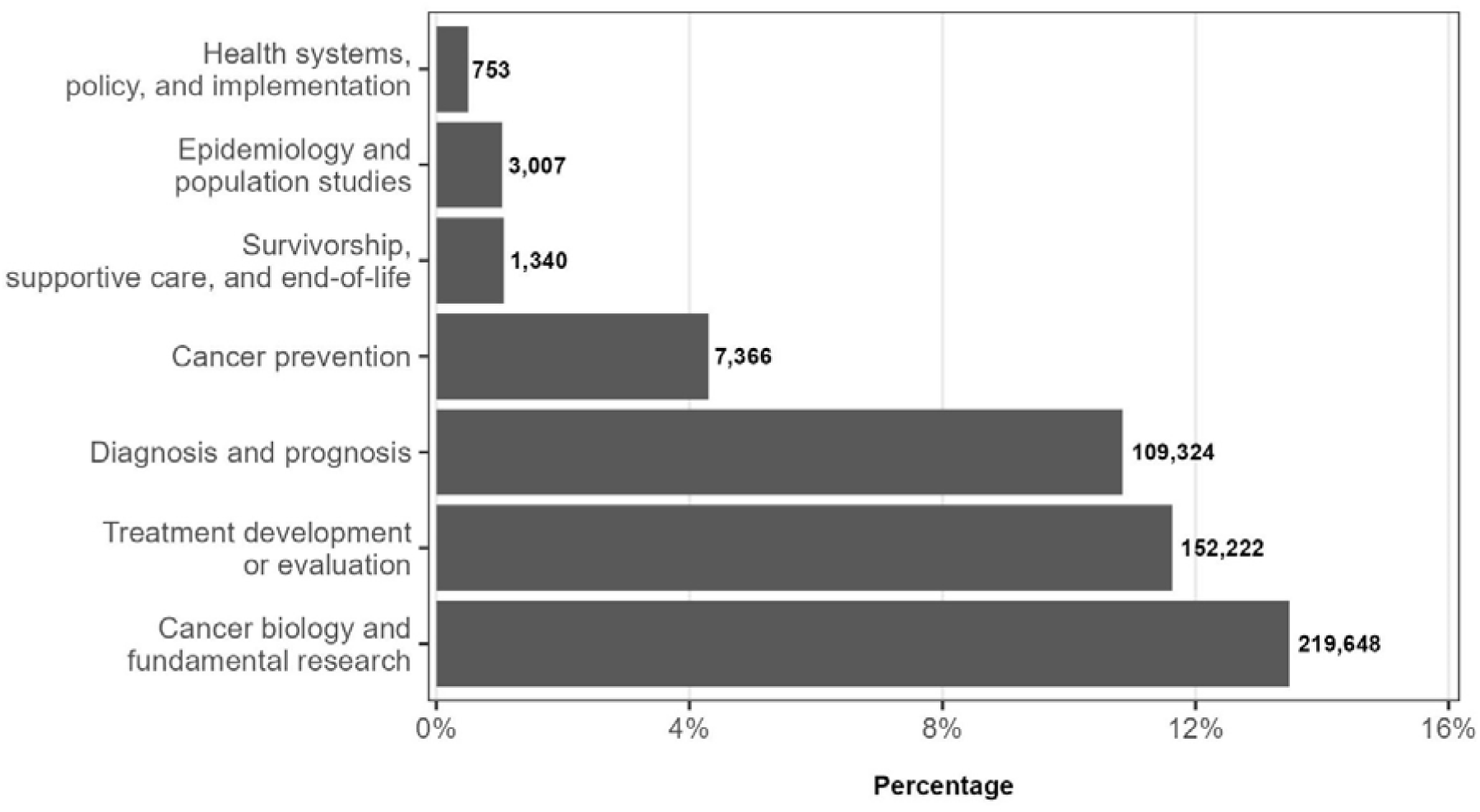
Percentage of cancer research papers within each research area, flagged because their titles and abstracts were similar to those of retracted paper mill papers. The number of flagged papers per area is shown next to the bars. As categories were assigned using a multi-labelling tool, each paper may appear in multiple categories, and the sum of all categories does not equal the total number of flagged papers. Percentages represent the proportion of papers flagged as similar to retracted paper mill papers within each research area, calculated as (flagged papers in area / total papers in area) × 100.

#### Flagged cancer research papers in high impact factor journals

The percentage of flagged papers in the top 10% of journals from the cancer research corpus by impact factor (Decile 1 or D1) shows a clear increase over time (*Figure 8*). While the percentage remained low in the early 2000s, a sustained increase occurred in the following years, reaching around 10% in 2022. The minimum impact factor required to be among the top 10% of journals for each year (cut-off impact factor) also increased from 3 in 1999 to 7 in 2021.

**Figure 8:**
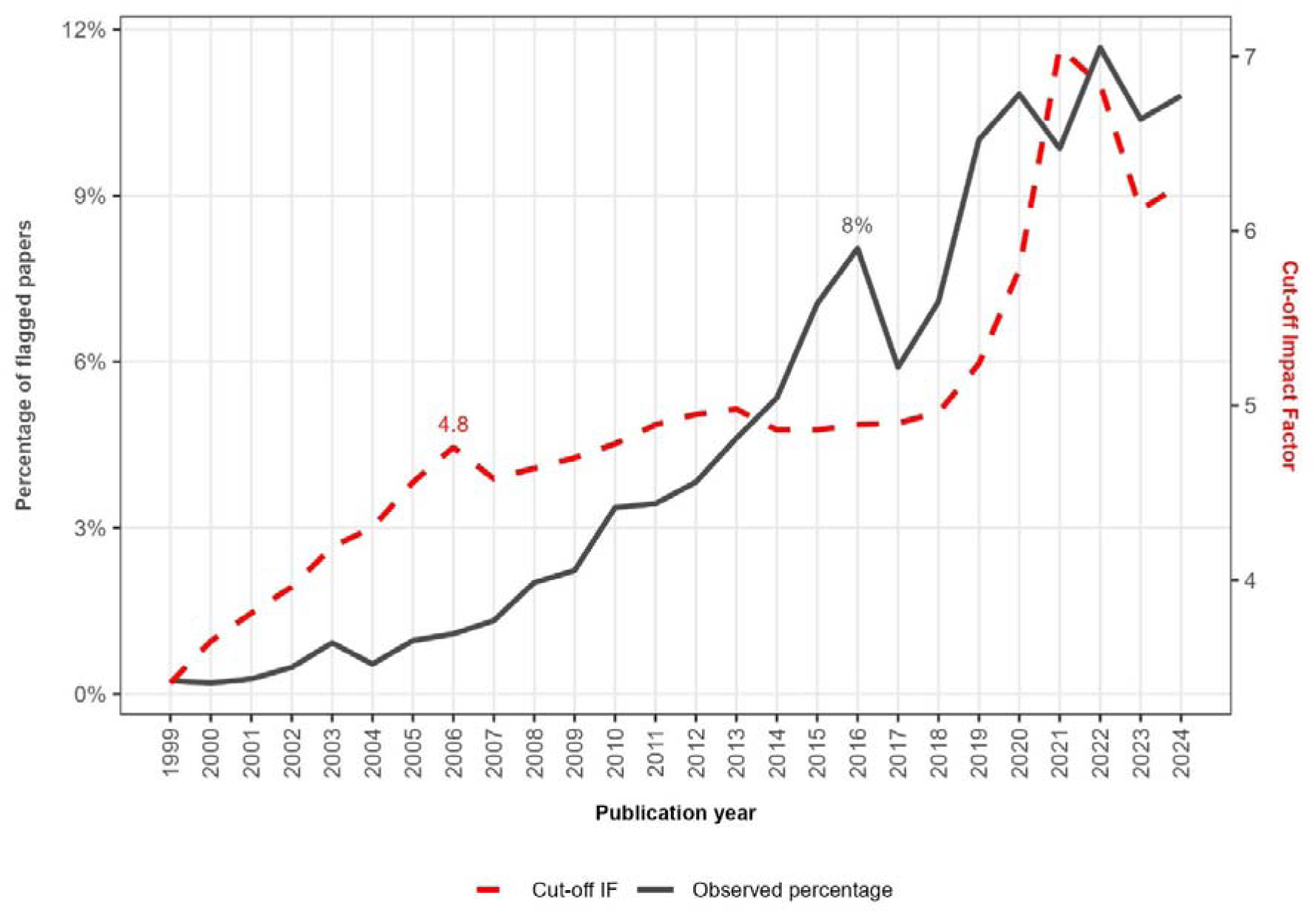
Percentage of cancer research papers flagged because their titles and abstracts were similar to those of retracted paper mill papers, in the top 10% of journals by impact factor, according to publication year. The minimum impact factor required to be among the top 10% of journals (cut-off impact factor) in each year is shown by a red dashed line. The 2006 cut-off impact factor (4.8) and the 2016 percentage of flagged papers (8%) are displayed as example figures. Please note that the Y-axis showing cut-off impact factor does not commence at zero.

## Discussion

We have fine-tuned a BERT machine learning model that achieves good accuracy in flagging suspected paper mill papers, using combined titles and abstracts from both Retraction Watch and image integrity experts’ datasets. The model also flagged many papers previously identified as containing incorrect nucleotide sequences or cell lines^42–44^, despite not having access to this information, demonstrating its ability to detect problematic papers without circular validation.

The model’s performance on Retraction Watch and integrity experts’ data supports the hypothesis that paper mills rely on textual templates to create the titles and abstracts of cancer research papers. While some of the text signals might be due to geographic writing styles, specific terminology, or area-specific research topics, the analysis of misclassified papers showed little systematic bias with a low number of false positives. Concerns about our model only identifying the writing style of Chinese authors are somewhat ameliorated by the high number of false *negatives* for authors from China, whereas we would be concerned about undue geographic discrimination if any country was over-represented in the false *positives*.

The model flagged almost 10% of cancer papers, a higher proportion than the previous 3% estimate of paper mill paper prevalence in biomedical research^3^, although those estimates were derived from different datasets, time periods, and detection criteria. The positive predictive value (PPV) depends on the true prevalence of paper mill papers, which remains unknown but is likely higher in cancer research^1^. Assuming a 10% true prevalence and the observed model performance (sensitivity and specificity), the PPV would be approximately 0.7, meaning that around 30% of flagged papers would represent false positives. This reinforces the need to not solely rely on the model prediction to conclude that a manuscript is from a paper mill.

Several indirect indicators support the validity of the model’s predictions: the exponential trend over time in flagged papers coincides with the known development of paper mill papers^2^. China as the leading country in terms of flagged papers is consistent with findings of paper mill paper origins^9,46,47^. Publishers with a high percentage of flagged papers in this study have been found to publish problematic papers, and some have been flagged as ‘*in conflict with academic rigour*’ by the Chinese government^47–49^. Flagged papers were more common in fundamental research which coincides with evidence from the literature^47^. Finally, suspected paper mill papers have already been identified in high-impact factor cancer journals^43^.

The exponential rise of flagged papers in *Figure 3* flattens after 2022. Three potential hypotheses can explain this phenomenon: the result of the publishers and research community fighting back against paper mills; a shift to new templates by paper mills, following the rise of AI; or the known delays in PubMed listing all papers for the most recent years. The relatively low number of flagged papers prior to 2010 may reflect the distribution of the training data, which primarily includes retracted paper mill papers published between 2013 and 2023, rather than indicating a near complete absence of such features during these years. This could also reflect the use of proxy labels rather than verified ground truth. Some unlabelled older papers may in fact be paper mill products, inflating apparent time trends and contributing to temporal drift.

The higher proportion of flagged papers in gastric and liver cancer research may partly be explained by the high prevalence of these cancers in China^50^. However their marked overrepresentation among misidentified cell lines – 25% and 15% of all such lines, respectively^51^ – is striking. Given that some misidentified cell lines, such as BGC-823 and BEL-7402, appear almost exclusively in publications from Chinese institutions^51^, this pattern may also reflect vulnerabilities exploited by paper mills, where popular research topics are targeted. It may also result from inertia, as early templates were reused and adapted repeatedly in these domains.

The rise in the percentage of flagged papers in Decile 1 journals suggests that paper mill papers are not just a low-impact journal problem. The concurrent increase in impact factors and the spread of flagged papers suggest that both phenomena may stem from the pressures of the *publish-or-perish* culture^52^. The increase of paper mills in high impact factor journals highlights an important limitation of using impact factors as proxies for research quality^43^.

Our training set of paper mill papers has limitations. The tag ‘*paper mill*’ in the Retraction Watch Database reflects only Retraction Watch staff’s interpretation of the publisher’s retraction notice. There is no uniformity in the way publishers investigate fraudulent papers and no standard way of interpreting the retraction notices; thus, the ‘*paper mill*’ qualification likely reflects varied levels of evidence. The papers listed online by research integrity experts include evidence of image manipulation, which can occur within settings beyond paper mills. Additionally, the experts may vary in their methods and transparency. Research on paper mills remains limited^1^, which could mean that the currently identified paper mill papers represent only a fraction of their actual prevalence in the scientific literature. Consequently, the model will likely not detect textual features associated with paper mill papers as a whole, but rather those represented in the training set.

The overrepresentation of authors from Chinese institutions among retracted papers suspected of originating from paper mills introduces a potential bias. Despite efforts to balance by language in the control set, a residual risk remains that the model may learn to associate linguistic patterns of Chinese scientific writing with paper mill content, rather than identifying features specific to fraudulent manuscripts. However, the model misclassification analysis shows few false positives, and an overrepresentation of Chinese papers among false negatives – which does not indicate systematic over-flagging. Furthermore, the paper mills’ country of origin can differ from the authors’ countries and Abalkina showed that a Russian paper mill sold publications to ‘*more than 800 scholars affiliated with more than 300 universities from at least 39 countries*’^16^. The misclassifications may instead reflect blind spots in the training data, where certain textual features in the wider population of paper mill papers were underrepresented or absent in our training data.

Additional sources of bias may stem from the composition of the control set. Controls were not randomly sampled from the broader cancer research literature to avoid including undetected paper mill papers. As a result, only a limited number of verified high-quality Chinese publications were included in the controls to minimise potential overlap with paper mill content. The assumption – supported by retraction data – that articles published in selected high impact journals or authored by Taiwanese, Swedish, Norwegian and Finnish research teams can serve as proxies for high-quality controls is open to criticism. While this strategy may enhance contrast between genuine and fabricated texts, it may also limit the model’s ability to detect more nuanced cases of fabrication. We also acknowledge that only limited human verification was conducted to assess the validity of the ground truth labels, and that potential misclassifications may have affected the model’s quality. Future research could manually verify a larger and more diverse set of Chinese publications to improve label accuracy.

The non-explainability of deep learning models prevents us from directly identifying the features captured by BERT. Flagged papers can include actual paper mill features; other features of misconduct; original work copied by paper mills; original work drawing inspiration from paper mill papers; and mistakes by the model. This research does not aim to directly identify paper mill papers or to accuse anyone of fraud, but rather to identify potentially problematic papers. The classifier is a probabilistic model, not a definitive arbiter of misconduct. As such, all flagged papers represent statistical predictions based on textual features and should be interpreted as signals requiring human judgment and further verification, not as confirmed cases of fraud.

Our model could be continuously improved by updating the training set with the latest confirmed paper mill papers. Since only titles and abstracts were used to train the model, incorporating full-text data or selected sections of the full-text has the potential to further enhance its performance. Future work could explore alternative, less computationally intensive training strategies, which might prove as effective as our fine-tuned BERT classifier, considering that we evaluated only a limited set of configurations. Experimenting with other aggregation strategies and post hoc calibration methods may also help to improve the robustness and interpretability of model predictions.

We expect the paper mills to react and innovate, as detection methods like ours threaten their income. The release of OpenAI’s ChatGPT-3.5 in 2022 and the rise of generative AI might further blur the boundaries between genuine and fabricated texts, rendering future automated detection of fraudulent features more challenging. While efforts to combat paper mills may evolve into an arms race^1^, this problem has reached an unacceptable scale. Inaction risks allowing paper mills to spread further, potentially compromising entire journals and publishers – as already seen in the case of Hindawi^5^.

Our model is currently integrated into the online submission systems of three journals from a major publisher and is being used to screen cancer-related manuscripts. To prevent unfair paper mill attributions, final decisions are always made by humans, using their expertise and a multi-tool detection approach to support decision-making. In fact, the predictions are intended to be informative and to precede a second phase of scrutiny. Thus, submissions are never rejected on the basis of the tool alone but only following serious integrity findings. Authors are not informed if their paper is flagged, in order to prevent paper mills from adapting their templates. This collaboration did not influence the writing of this manuscript and no data was collected throughout this pilot testing phase.

In conclusion, this study demonstrates that using machine learning to identify papers resembling retracted paper mill papers from titles and abstracts is both feasible and effective. Our findings reveal concerning trends in cancer research publishing. The rising percentage of flagged papers indicates that paper mills have grown in ambition and now target higher impact factor journals, highlighting the need for vigilance among journals, reviewers, and researchers. While our model has clear limitations, it provides useful insights and highlights the need for collective awareness to curb the spread of paper mill publications.

## Author contributions

Conceptualisation: AGB and JAB. Methodology: BS, AGB, DC and JAB. Code development and execution: BS. Analysis: BS, AGB, DC and JAB. Writing: BS and AGB. Reviewing and editing: BS, AGB, DC and JAB. Funding acquisition: JAB and AGB. Supervision: AGB, JAB and DC. The work reported in the paper has been performed by the authors only.

## Funding

This study was funded by the National Health and Medical Research Council (NHMRC), Ideas Grant no. 2029249: ‘*Problematic Articles and Literature Reviews in Molecular Cancer Research*’

## Ethics statement

Not applicable.

## Data and code availability statement

All data used in this study are publicly available: PubMed annual XML datasets (https://ftp.ncbi.nlm.nih.gov/pubmed/baseline/), Retraction Watch database^27^, and the integrity experts’ dataset^32^. The fine-tuned models and code developed in this study are not publicly disclosed to prevent potential misuse by individuals seeking to evade fraud detection. The list of flagged PMIDs is available upon reasonable request.

## Acknowledgement

The authors thank Associate Professor Nathalie Bock for her valuable review of this manuscript. We acknowledge the members of the high-performance computing environment teams of Aqua (Queensland University of Technology, QUT), Bunya (Queensland Cyber Infrastructure Foundation, QCIF, on behalf of the University of Queensland, UQ) and Tesla (*Institut de recherche mathématique de Rennes*, IRMAR, France) for granting access to their resources and for their technical support. We also acknowledge the teams at PubMed, Retraction Watch, SCImago, and the contributors from PubPeer, especially anonymous user *Hoya Camphorifolia*, for sharing their data and insights.

## Competing interest statement

The authors declare no competing interests.

## Notes

### Competing Interest Statement

The authors have declared no competing interest.

### Summary of Updates

1) The introduction was revised to provide more detail on the study's motivation and main hypothesis. 2) The methods section was clarified, and a new summary table was added. 3) The model misclassification analysis was corrected and clarified; a new table was added, and the Chi-square table was simplified. 4) The discussion was improved and reorganised. 5) New supplementary files were added to provide additional information.

