## Supplementary file 1 for "Revealing the Paper Mill Iceberg: AI-Based Screening of Cancer Research Publications"

**Supplementary File 1: Methods**

**Affiliation-Based Country Indexing**

The country of origin for each cancer research paper was inferred from the first author’s affiliation. Affiliation addresses were processed to search for country names or common acronyms such as UK or UAE, using the ISO 3166-1 standard via the *pycountry* package^1^. For 432,145 papers, the country of origin could not be identified due to missing or incomplete affiliation information. In cases where the country name was missing but the rest of the affiliation was present, the addresses were passed through Google FLAN-T5-small^2^ to infer the most likely country. Ultimately, 41,007 papers lacked any affiliation data, and for 89,446 others, the country could not be determined from incomplete affiliation information – resulting in a total of 130,453 papers labelled as 'No Data'. These data were excluded from country-related visualisations.

**Publisher Identification**

Using the journal ISSNs linked to papers from the cancer research corpus, the publishers were identified using the SCImago database^3^. A total of 137,361 (5%) papers could not be associated with a publisher via their ISSN, as the corresponding ISSNs were not indexed in SCImago.

Over the years 1999–2024, 1,822 publishers were identified as having contributed to the publication of the remaining 2,510,110 papers that included ISSNs. To simplify the analysis, only major publishers were retained in the dataset: publishers were ranked in descending order using the number of cancer-related publications, and those cumulatively accounting for 95% of cancer papers were retained. This selection process resulted in the retention of 212 out of 1,822 publishers, covering 2,384,049 papers (95%). The remaining 1,610 publishers each contributed to 126,061 papers over the years 1999–2024.

The 212 selected publishers were assigned to their parent companies. For example, *Taylor & Francis*, *Landes Bioscience*, and *Informa Healthcare* were grouped under *Informa*. Following this grouping, 136 distinct publishers were retained. Other publishers, as well as ISSNs for which no publisher could be identified, were assigned to the category 'Other', representing 263,422 papers (10%). These data were excluded from publisher-related visualisations.

**Cancer-type identification**

The type of cancer investigated in the papers was determined using AI labelling. Cancer type classification was based on cancer prevalence, adapted from the International Agency for Research on Cancer (IARC)^4^ data (Box S1). Other cancer types were classified as 'unspecified cancer'. Data corresponding to 'unspecified cancer' were excluded from the visualisations.

**Box S1: Cancer type classification.**

1. breast cancer, 2. lung cancer, 3. colorectal cancer, 4. prostate cancer, 5. gastric cancer,
6. pancreatic cancer, 7. liver cancer, 8. ovarian cancer, 9. cervical cancer, 10. uterine cancer, 11. kidney cancer, 12. bladder cancer, 13. brain cancer, 14. skin cancer, 15. thyroid cancer, 16. blood cancer, 17. head and neck cancer, 18. oesophageal cancer, 19. bone cancer, and
20. unspecified cancer

Gastric cancer includes stomach and biliary tract cancers; brain cancer includes brain and central nervous system (CNS) cancers; blood cancer includes leukemia, lymphoma, and multiple myeloma; and head and neck cancer includes oral, laryngeal, pharyngeal, nasal, and salivary gland cancers.

A randomly selected subset of 15,000 papers from the cancer research corpus were pre-labelled using gpt-4o-mini from the OpenAI API on the title, abstract and MeSH terms of each paper. The prompt is displayed in Box S1.1 (Temperature = 0). A sample of 100 papers was double-annotated by a human reader (AB) and compared to GPT-4o-mini’s annotations, resulting in an unweighted Cohen’s Kappa of 0.81 (95% CI 0.72 to 0.89). All 15,000 labelled papers were used to train a BioBERT^5^ model for a text-classification task to label the whole cancer research corpus (with accuracy = 0.95, F1-score micro = 0.94, F1-score macro = 0.92).

**Box S1.1: Prompt for cancer type classification**

""" You will be given a medical abstract or paragraph.

Your task is to classify the specific type of cancer being discussed. Choose only one label from the list below.

Read the text carefully and identify whether a clearly defined cancer type is mentioned based on biological, anatomical or clinical context.

- If the text explicitly mentions a cancer that matches one of the listed types, select that type.

- If multiple cancer types are mentioned select the one that is most central to the study, based on experimental models (e. g. cell lines, animal models), patient population or focus of the abstract.

- If the text refers to a cancer but the type is unclear, general, or not on the list, select unspecified cancer.

Respond with exactly one label , using the list below:

1. breast cancer

2. lung cancer

3. colorectal cancer        #Includes colon and rectal cancer

4. prostate cancer

5. gastric cancer    # includes stomach and biliary tract cancers

6. pancreatic cancer

7. liver cancer

8. ovarian cancer

9. cervical cancer

10. uterine cancer

11.kidney cancer

12. bladder cancer

13. brain cancer  # Includes brain and central nervous system (CNS) cancers

14. skin cancer

15. thyroid cancer

16. blood cancer  # Includes leukemia, lymphoma and multiple myeloma

17. head and neck cancer # Include oral, laryngeal, pharyngeal, nasal and salivary gland cancers

18. oesophageal cancer

19. bone cancer

20. unspecified cancer

Now classify the following text:"""

**Cancer research classification**

As research objectives are often diverse and cannot always be captured by a single label, a multi-label classification approach was adopted to categorise the cancer research objectives using AI-based labelling. The classification scheme was built around a set of key research aims relevant to cancer research adapted from NCI – National Cancer Institute^6^ research areas (see Box S2). Importantly, clinical trials were excluded from this classification, as such papers were filtered out of the cancer research corpus (due to our focus on preclinical research).

**Box S2: Cancer research classification**

1. Cancer biology and fundamental research, 2. Treatment development or evaluation, 3. Diagnosis and prognosis, 4. Prevention, 5. Survivorship, supportive care, and end-of-life, 6. Epidemiology and population studies, 7. Health systems, policy, and implementation

A representative subset of 12,000 papers was randomly sampled from the cancer research corpus and pre-labelled using the GPT-4o-mini model from the OpenAI API, based on each paper’s title, abstract, and MeSH terms. The prompt is displayed in Box S2.1 (Temperature = 0). A sample of 100 papers was double-annotated by a human reader (AB) and compared to GPT-4o-mini’s annotations. We computed Cohen’s Kappa for each class: 1 = 0.74, 2 = 0.56, 3 = 0.57, 4 = 0.86, 5 = 0.65, 6 = 0.70, and 7 = 0.64. The macro-averaged Kappa (unweighted mean across classes) was 0.68, and the micro-averaged Kappa (weighted by class frequency) was 0.66. All 12,000 annotated papers were then used to fine-tune a BioBERT model for a multi-label classification task, enabling automated labelling of the entire corpus (with multi-label accuracy = 0.7, F1-score micro = 0.9, F1-score macro = 0.87).

**Box S2.1: Prompt for cancer research classification**

""" You will be given a medical abstract or paragraph.

Your task is to classify the main objectives of the cancer research study described in the text.

Pick one or more categories from the list below that match the goals of the study, not the methods used, data collected or secondary details.

Respond with the list of codes that apply, separated by commas (BIO, THER).

Categories:

-BIO -Understanding cancer biology

- PREV - Prevention

- DIAG - Diagnosis and prognosis

- THER -Treatment development or evaluation

-IMPL - Health systems, policy, and implementation

- SURV - Survivorship, supportive care and end-of-life

- EPID - Epidemiology and population studies

Choose all codes that apply based on the primary objectives of the study.

Now classify the following text:"""

**SCImago Journal Impact Factor retrieval**

The SCImago Journal Impact Factor (SJIF) was retrieved for each journal from the SCImago database from 1999 to 2024. An SJIF value was available for 8,701 journals, while the remaining 2,931 journals were either not found or not indexed in SCImago. Each paper was matched with the SJIF corresponding to journal and year of publication. As a result, 2,469,177 papers (93%) were assigned an SJIF value, while 178,294 papers (7%) were labelled as 'no SJIF'.

**Hyperparameter optimisation**

The search space was pre-restricted through empirical testing, allowing the batch size to 32, as this setting yielded better performance. The number of training epochs was fixed at 10 to allow sufficient training time. The evaluation criterion was set to the evaluation loss to ensure greater model stability and prediction accuracy. Only the learning rate, weight decay, warmup ratio and scheduler type were optimised. Please note that this optimisation scheme only applies to the BERT model used for paper mill paper classification.

An initial hyperparameter search consisting of 100 trials was conducted using a guided exploratory strategy with random parameter combinations to identify promising configurations. The learning rate was searched within the range of 5e-6 to 5e-5, weight decay within 0.005 to 0.3, scheduler type between linear and cosine and the warmup ratio between 0% and 25% of the total training steps.

This was followed by a second, more focused adaptive search consisting of 50 trials to further refine the best-performing configuration. The top 10% of trials (based on evaluation loss) from the initial search were used to guide this step, with the goal of selecting the best-performing and most stable configuration. The learning rate search space was narrowed to the 1e-5 to 2e-5 range, and the warmup ratio to the 0.015 to 0.020 range. The weight decay showed an optimal range between 0.02 and 0.03. The scheduler type was fixed to cosine, with 'cosine with restarts' also evaluated. Final optimization results indicated the following parameters: a learning rate of 1.4e-5, a weight decay of 0.025, 15% warm-up steps, and a basic cosine scheduler type.

The final BERT model was trained for 2 hours using the optimised and fixed parameters and the 2100-training step checkpoint was selected (2 epochs), and its weights were merged with the initial model. After testing, the mean-over-probabilities method was preferred over logistic regression as the aggregation method, since the latter achieved similar performance (logistic regression: Accuracy = 0.91, Sensitivity = 0.92, and Specificity = 0.91). The decision threshold for flagging suspect papers was set at a probability of 0.6019, based on ROC curve optimisation. Large scale inferences on the cancer research corpus were made using a Tesla V100S-PCIE-32GB GPU over approximately 12 hours.

3.	SCImago. SJR — SCImago Journal & Country Rank. http://www.scimagojr.com.

4.	International Agency for Research on Cancer (IARC). https://gco.iarc.who.int/en.

5.	Lee, J. *et al.* BioBERT: A pre-trained biomedical language representation model for biomedical text mining. *Bioinformatics* **36**, 1234–1240 (2020).

6.	National Cancer Institute (NCI). https://www.cancer.gov/.
