## Supplementary file 2 for "Revealing the Paper Mill Iceberg: AI-Based Screening of Cancer Research Publications"

**Supplementary File 2: Cancer research corpus breakdown**

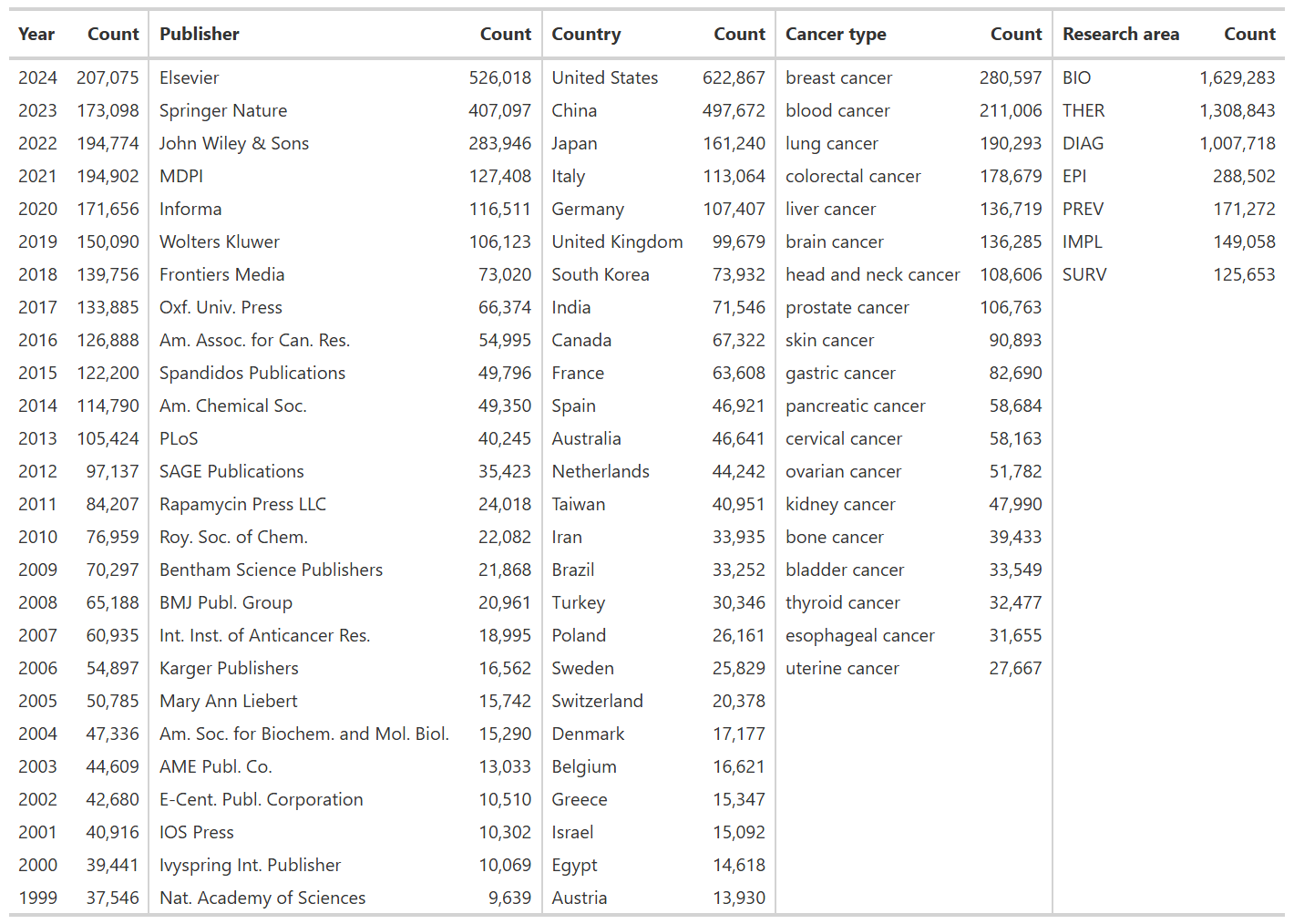
*Table S2: Summary of the cancer research corpus. The years have been displayed in descending chronological order and the number of rows has been set to the number of years (26). The other columns are displayed in descending order according to their counts.*
