## Supplementary file 3 for "Revealing the Paper Mill Iceberg: AI-Based Screening of Cancer Research Publications"

**Supplementary File 3: Paper mill data**

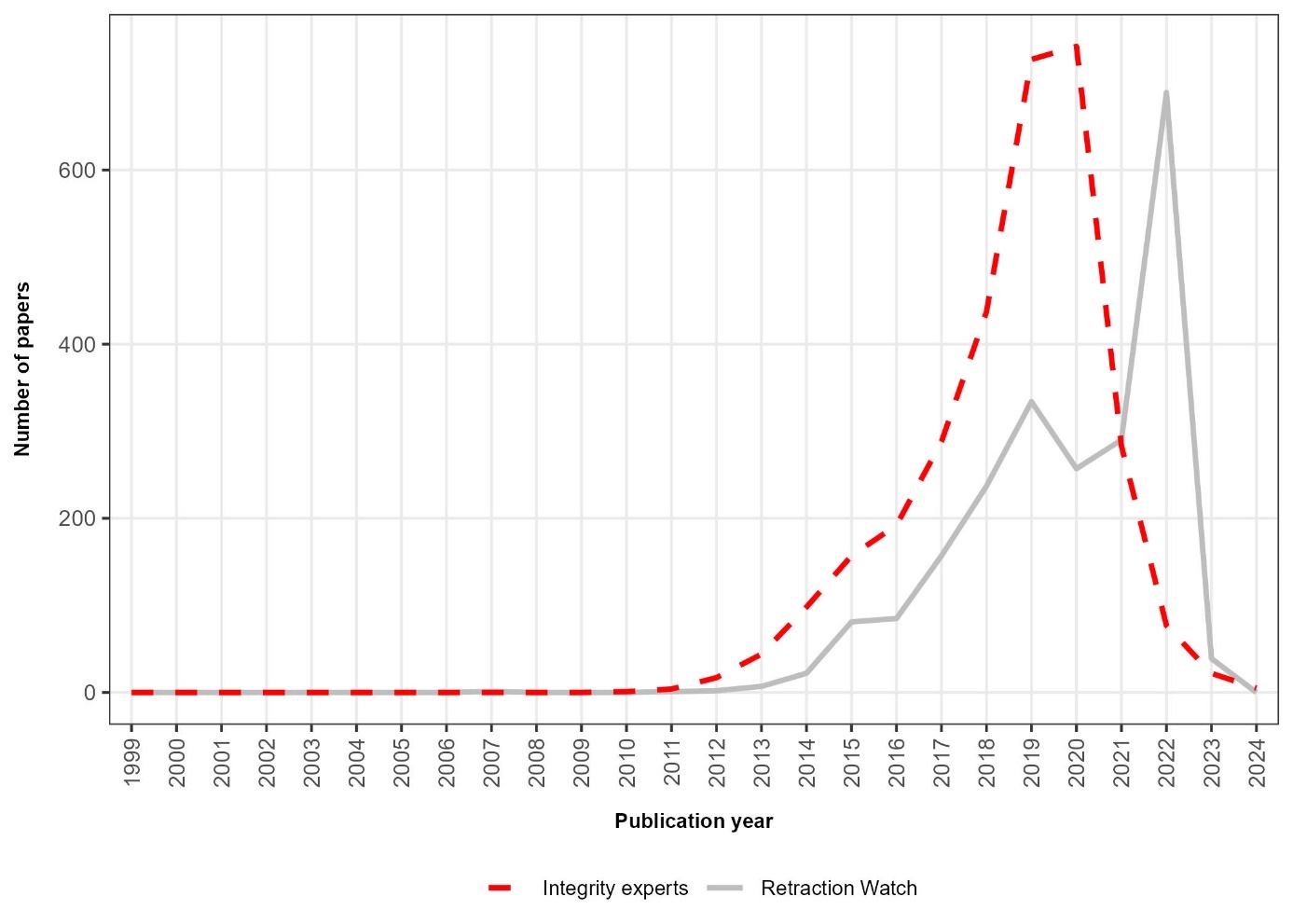

*Figure S3.1: Distribution of retracted paper mill papers in the Retraction Watch (2,202 papers) and suspected paper mill papers from the integrity experts’ set (3,094 papers). The Retraction Watch set is a solid grey line while the experts’ set is a red dashed line. All papers present in both datasets (~1,000 papers) were removed from the integrity experts’ set. The Retraction Watch dataset was used for training, validation, and testing, while the integrity experts’ dataset was used exclusively for testing.*

*Table S3.1:* *Ten most frequent unigrams, bigrams, and trigrams found in the titles of retracted paper mill papers indexed by Retraction Watch.*

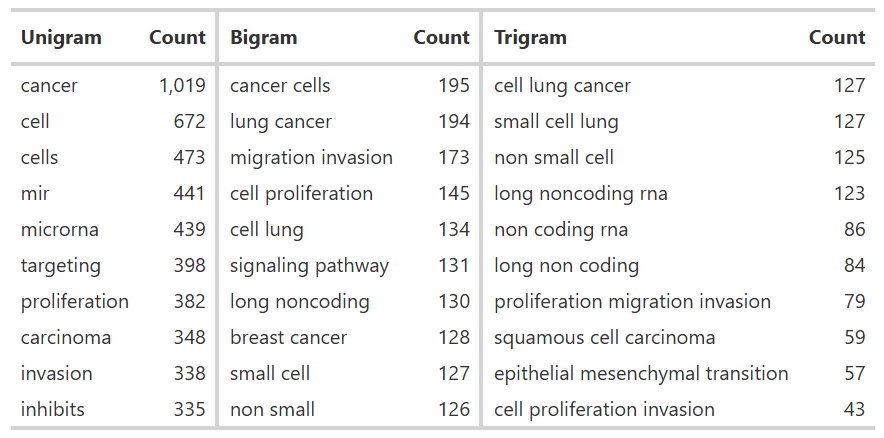

*Table S3.2: Five most frequent publishers, first author country of affiliation, cancer types investigated and main research axes among retracted paper mill papers indexed by Retraction Watch. Entries with no country data (n = 34) and unspecified cancer types (n = 229) were excluded from the table. Research area labels: BIO - Cancer biology and fundamental research; THER - Treatment development or evaluation; DIAG - Diagnosis and prognosis; SURV - Survivorship, supportive care, and end-of-life; and PREV - Cancer prevention.*

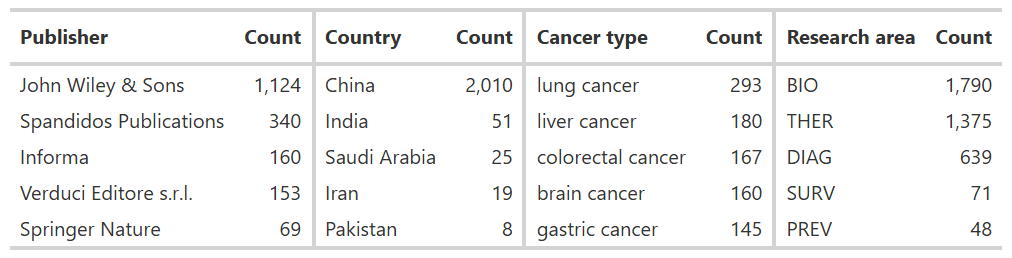

*Table S3.3:* *Ten most frequent unigrams, bigrams, and trigrams found in the titles of suspected paper mill papers listed in the integrity experts’ dataset.*

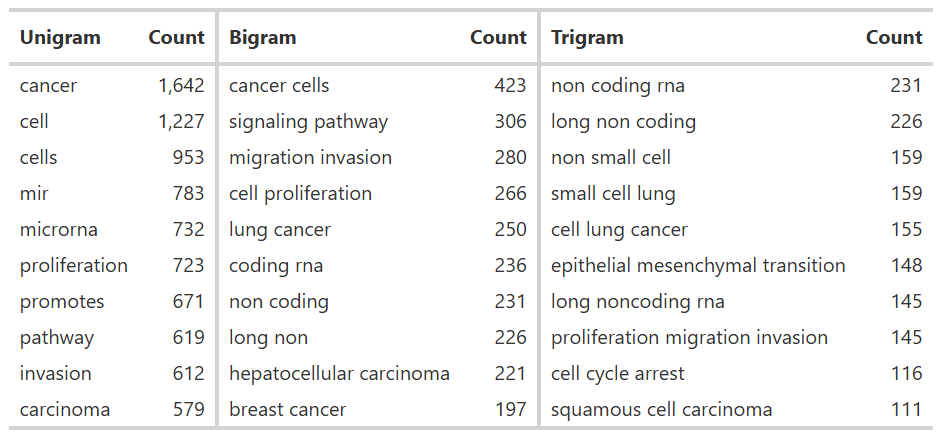

*Table S3.4: Five most frequent publishers, first authors country of affiliation, cancer types investigated and main research areas among suspected paper mill papers listed in the integrity experts’ dataset. Research area labels: BIO - Cancer biology and fundamental research; THER - Treatment development or evaluation; DIAG - Diagnosis and prognosis; SURV - Survivorship, supportive care, and end-of-life; and PREV - Cancer prevention.*

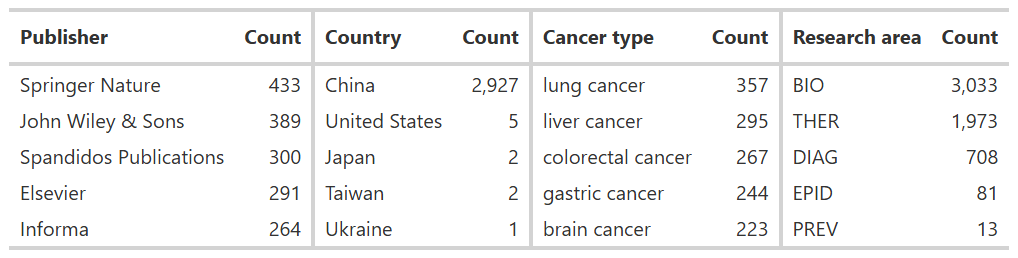
