## Supplementary file 4 for "Revealing the Paper Mill Iceberg: AI-Based Screening of Cancer Research Publications"

**Supplementary File 4: Model pre-assessment**

We have conducted preliminary experiments with BERT and other BERT-based models, including RoBERTa, BioBERT, PubMedBERT, Longformer, and Clinical Longformer. These alternatives were selected to assess whether biomedical-specific pretraining or extended input capacity (up to 4,096 tokens, compared to BERT’s 512-token input limit) could enhance classification performance. While PubMedBERT and BioBERT slightly outperformed BERT in our empirical experiments, we did not consider these differences as important given BERT’s already high performance (*Table S4*). As neither domain-specific pretraining nor the ability to process longer text sequences provided a clear advantage over BERT and given the potential risk of data leakage in domain-specific models, we chose to retain BERT.

BERT, RoBERTa, BioBERT, PubMedBERT, Longformer, and Clinical Longformer were not trained and assessed on the same data as those used in the manuscript. Although the overall methodology was identical for the origin of cases and the selection of controls, early testing was conducted in June 2024. Approximately 1,400 paper mill papers were used for training, while the external validation set comprised the remaining 200 Retraction Watch papers and 2,000 papers identified by Integrity Experts. Paper mill papers were matched with an equal number of controls (n = 3,400), selected from Chinese (5%), Taiwanese (28%), Finnish (11%), Swedish (11%), and Norwegian papers (11%), as well as papers published in high-impact journals (33%). Accuracy, sensitivity, and specificity were derived from the confusion matrix using a probability threshold to prioritise specificity and minimise false positives.

*Table S4: Initial empirical comparison of six models. Preliminary experiments compared BERT, RoBERTa, BioBERT, PubMedBERT, Longformer, and Clinical Longformer to assess the impact of biomedical-specific pretraining and extended input capacity on classification performance, using Retraction Watch and integrity experts’ data as June 2024. All metrics show similar predictive performance.*

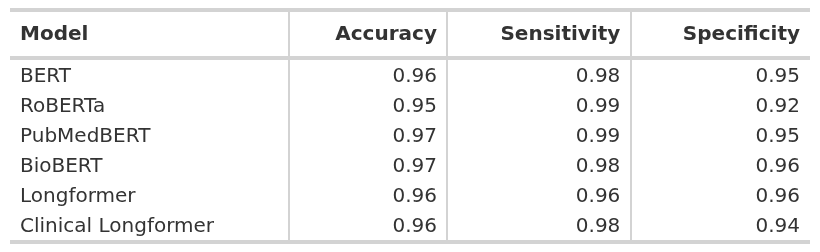
