## Supplementary file 5 for "Revealing the Paper Mill Iceberg: AI-Based Screening of Cancer Research Publications"

**Supplementary File 5: Examples of papers flagged with high probability**

All three papers presented in Supplementary File 5 are retracted papers tagged as '*Paper Mill*' in the Retraction Watch database. They were selected from the test set and were not seen by the model during training.

**Retracted paper A**

Model-predicted paper mill origin probability: **0.99**

**Title**: Long non-coding RNA SNHG14 contributes to gastric cancer development through targeting miR-145/SOX9 axis

**Abstract**: This study aimed to elucidate the roles of long non-coding RNA SNHG14 in gastric cancer development. LncRNA SNHG14 was markedly up-regulated in gastric cancer tissues and cells. Knockdown of SNHG14 significantly inhibited SGC-7901 cell viability, migration, invasion, and promoted cell apoptosis. In addition, miR-145 was negatively regulated by SNHG14 and the effects of SNHG14 knockdown on cell viability, apoptosis, migration, invasion, and the expression of apoptosis-related proteins and EMT-markers were reversed by inhibition of miR-145 at the same time. Furthermore, SOX9 was verified as a functional target of miR-145, and miR-145 regulated tumor malignant behaviors through regulating SOX9. Besides, knockdown of SNHG14 inhibited the expression of p-PI3 K, p-AKT, and p-mTOR and promoted PTEN expression, where miR-145 inhibition had opposite effects. Moreover, the activated PI3 K/AKT/mTOR pathway caused by miR-145 inhibition was counteracted after knockdown of SOX9. Our findings indicate that up-regulation of lncRNA SNHG14 may contribute to gastric cancer development via targeting miR-145/SOX9 axis and involving in PI3 K/AKT/mTOR pathway. SNHG14-miR-145/SOX9 axis may be a promising therapeutic strategy for gastric cancer treatment.

**Retracted paper B**

Model-predicted paper mill origin probability: **0.98**

**Title**: miR-451a Inhibits the Growth and Invasion of Osteosarcoma via Targeting TRIM66.

**Abstract**: The importance of microRNAs in regulating osteosarcoma development has been studied in recent years. However, the function of microRNA-451a in osteosarcoma growth is rarely investigated. Here, we explored the expression of microRNA-451a in osteosarcoma cell lines. Bioinformatic software, luciferase activity reporter assay, and Western blot were conducted to determine the association between microRNA-451a and tripartite motif-containing 66. Cell Counting Kit-8 assay and transwell assay were used to explore the regulatory effects of microRNA-451a on osteosarcoma cells. Moreover, we explored whether microRNA-451a modulates osteosarcoma cell biological activity by regulating tripartite motif-containing 66. The expression of microRNA-451a was found to be downregulated in osteosarcoma and negatively regulated the expression of tripartite motif-containing 66. Tripartite motif-containing 66 was further validated as a target of microRNA-451a. MicroRNA-451a inhibits the growth and invasion of osteosarcoma cell lines through targeting tripartite motif-containing 66. The miR-451a targets tripartite motif-containing 66 may provide novel therapeutic targets for the treatment of osteosarcoma.

**Retracted paper C**

Model-predicted paper mill origin probability: **0.98**

**Title**: SLCO4A1-AS1 mediates pancreatic cancer development via miR-4673/KIF21B axis.

**Abstract**: In this study, we intended to figure out the biological significance of long non-coding RNAs (lncRNAs) solute carrier organic anion transporter family member 4A1 antisense RNA 1 (SLCO4A1-AS1) in pancreatic cancer (PC). Cell counting kit-8, colony formation, wound healing, transwell, and flow cytometry experiments were performed to reveal how SLCO4A1-AS1 influences PC cell proliferation, migration, invasion, and apoptosis. Thereafter, bioinformatics analysis, RNA immunoprecipitation assay, luciferase reporter assay, and RNA pull-down assay were applied for determining the binding sites and binding capacities between SLCO4A1-AS1 and miR-4673 or kinesin family member 21B (KIF21B) and miR-4673. The results depicted that SLCO4A1-AS1 was upregulated in PC, and SLCO4A1-AS1 knockdown suppressed PC cell growth, migration, invasion, and induced cell apoptosis. Furthermore, SLCO4A1-AS1 was verified to modulate the expression of KIF21B by binding with miR-4673. SLCO4A1-AS1 exerted an oncogenic function in PC. The overexpression of SLCO4A1-AS1 aggravated the malignant behaviors of PC via the upregulation of KIF21B by sponging miR-4673. Our findings revealed a novel molecular mechanism mediated by SLCO4A1-AS1, which might play a significant role in modulating the biological processes of PC.
