## Supplementary file 6 for "Revealing the Paper Mill Iceberg: AI-Based Screening of Cancer Research Publications"

**Supplementary File 6: Supplementary analysis of model misclassifications**

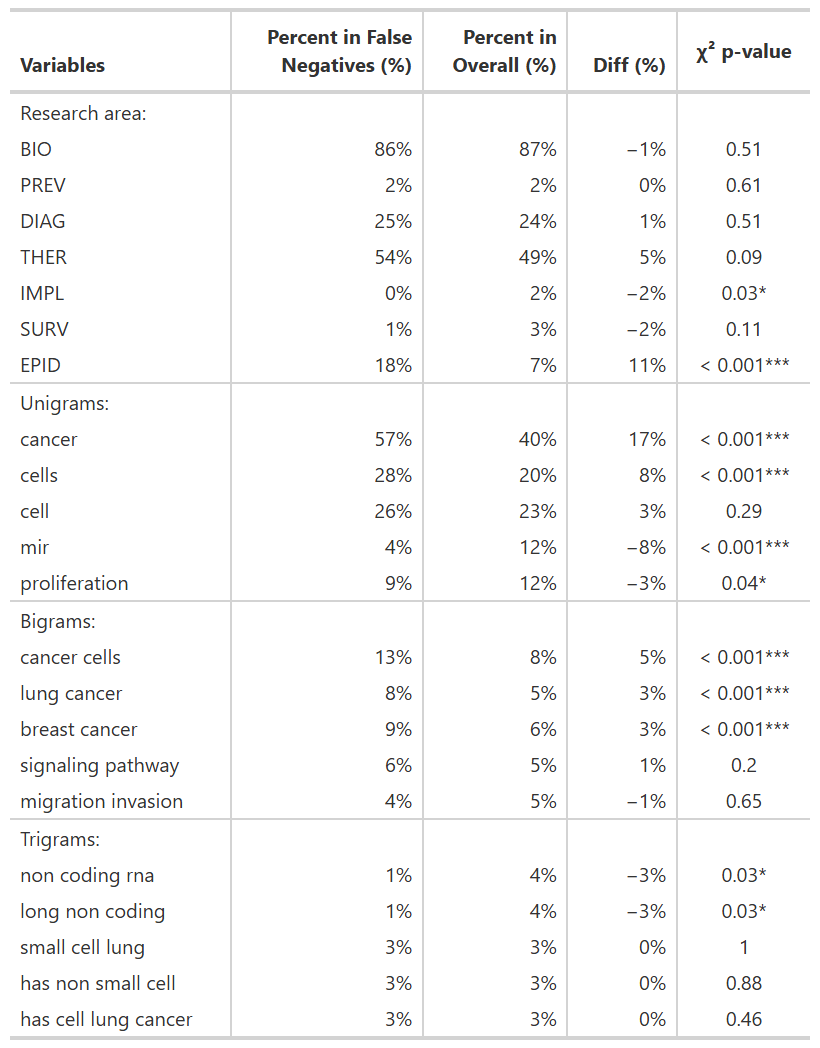
*Table S6. Characteristics of false negatives (n = 433) by research area, and title n-grams (unigrams, bigrams and trigrams). A Pearson’s chi-squared test of independence was carried out for each category of the multi-label variables (multiple testing). P-values were corrected with the Benjamini-Hochberg method. The percentage is shown both within the false negatives and in the overall pooled validation dataset (n = 6,745). The difference represents false negatives minus overall. Research area codes: BIO – Cancer biology and fundamental research; THER – treatment development or evaluation; DIAG – Diagnosis and prognosis; EPID – epidemiology and population studies; PREV – Prevention; SURV – Survivorship, supportive care, and end-of-life; IMPL – Health systems, policy, and implementation.*
